# MenDEL: automated search of BAC sets covering long DNA regions of interest

**DOI:** 10.1101/2022.06.26.496179

**Authors:** Sergei German, Sudarshan Pinglay, Brendan Camellato, David Fenyö, Jef D. Boeke

## Abstract

**Motivation:** Synthetic genomics as a field seeks to synthesize large regions of genomes from the ground up. Such large-scale projects, especially in complex genomes can rely on pre-existing BAC (Bacterial Artificial Chromosome) libraries as starting material to reduce cost. However, choosing BACs that cover long DNA regions, especially those that require many BACs, is a manual, idiosyncratic, time consuming, and error prone process. Automating this work would make the assembly of large DNA constructs more efficient.

**Results:** We have developed MenDEL – a web-based DNA design application, that provides efficient tools for finding BACs that cover long regions of interest and allow for sorting results based on multiple user defined criteria - total length, number of BACs, longest BAC. etc. In addition, it enables the user to find a combination of BACs from pre-existing libraries that cover a region of interest not found in any single BAC.

**Availability and Implementation:** MenDEL application is available to registered users at https://mendel-isg.nyumc.org, Java code used in the application to find BAC sets is available at https://github.com/MendelProject/BACFinder

## 1. Introduction

The construction of synthetic DNA assemblies is a central process in molecular and synthetic biology. It has recently become possible to synthesize large DNA constructs (>100 kilobases) and even entire chromosomes for the study of fundamental biological processes as well as to increase the capacity for genetic engineering (1). As these ‘synthetic genomics’ efforts move towards more complex organisms with larger genomes, the cost of chemically synthesizing any substantial fraction remains challenging.

Bacterial Artificial Chromosome (BACs) libraries from many species were constructed during early genome sequencing efforts and are publicly available in repositories - e.g. BACPAC Resources Center (BPRC). These libraries often include thousands of clones, each containing inserts >100kb and may provide a cost-effective starting material for larger synthetic genomics efforts. Indeed, methods have been developed to combine multiple BACs into larger constructs in both yeast and bacteria.(2)

Despite the promise of BACs as cheap sources for synthetic genomics, it is currently challenging to choose BACs that correspond to a region of interest in any genome. This is especially true for large regions that require more than a single BAC to cover the span. When multiple sets of BACs from multiple libraries could be potentially used for building the assembly, choosing the optimal set becomes even more challenging. While some information on the genomic content and position of a limited number of libraries is available in public databases (NCBI, UCSC), the manual identification of BACs corresponding to a region of interest is a cumbersome and error-prone process. (Figure 1) Further, manual design does not scale to the full extent of complex genomes. In this work, we present an automated and user-friendly solution to the BAC design problem.

**Figure 1.**
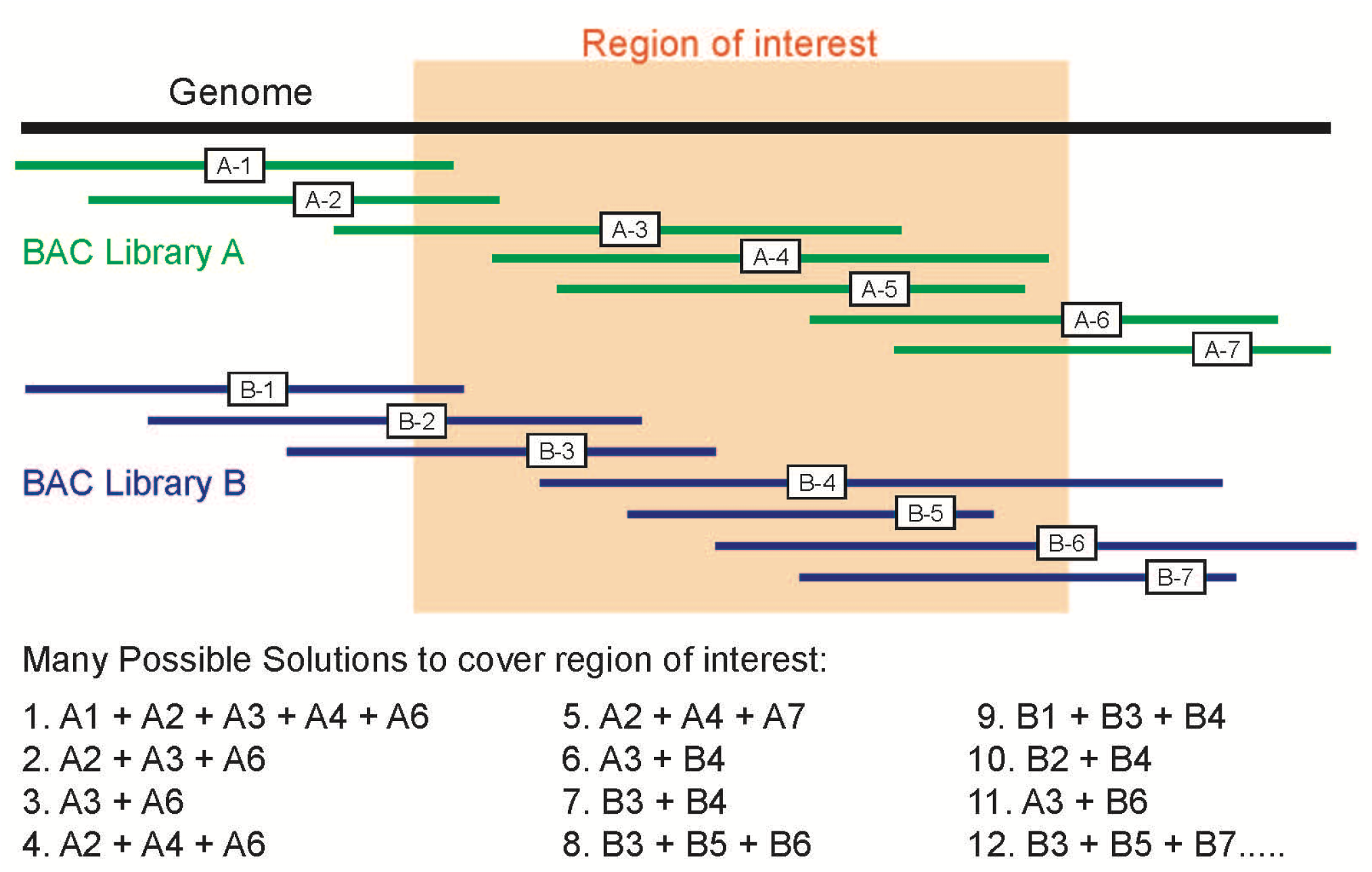
Problem of finding BAC solutions from multiple libraries for a given region of interest.

We call a set of BACs that cover the region of interest a “BAC solution”. We propose a method that allows us to automate and quantify the search of possible BAC solutions from multiple libraries. When the BAC solutions are found, we provide users with ways to both visualize solutions vis-a-vis each other and the region of interest itself. We also provide users with ways to filter and sort BAC solutions based on a number of characteristics - library, number of BACs in the solutions, total length of the solutions, length of longest BAC, etc.

## 2. Implementation

### 2.1 Data structures, algorithms, and class hierarchy

We interpret a set of BAC solutions as branches of a N-ary tree (3,4) Each node of the tree (Fig. 2) contains a reference to a BAC object and up to N references to the children nodes. For each available library we build a separate tree. The root of the tree has *null* reference to a BAC object, and up to N references to children nodes.

**Figure 2.**
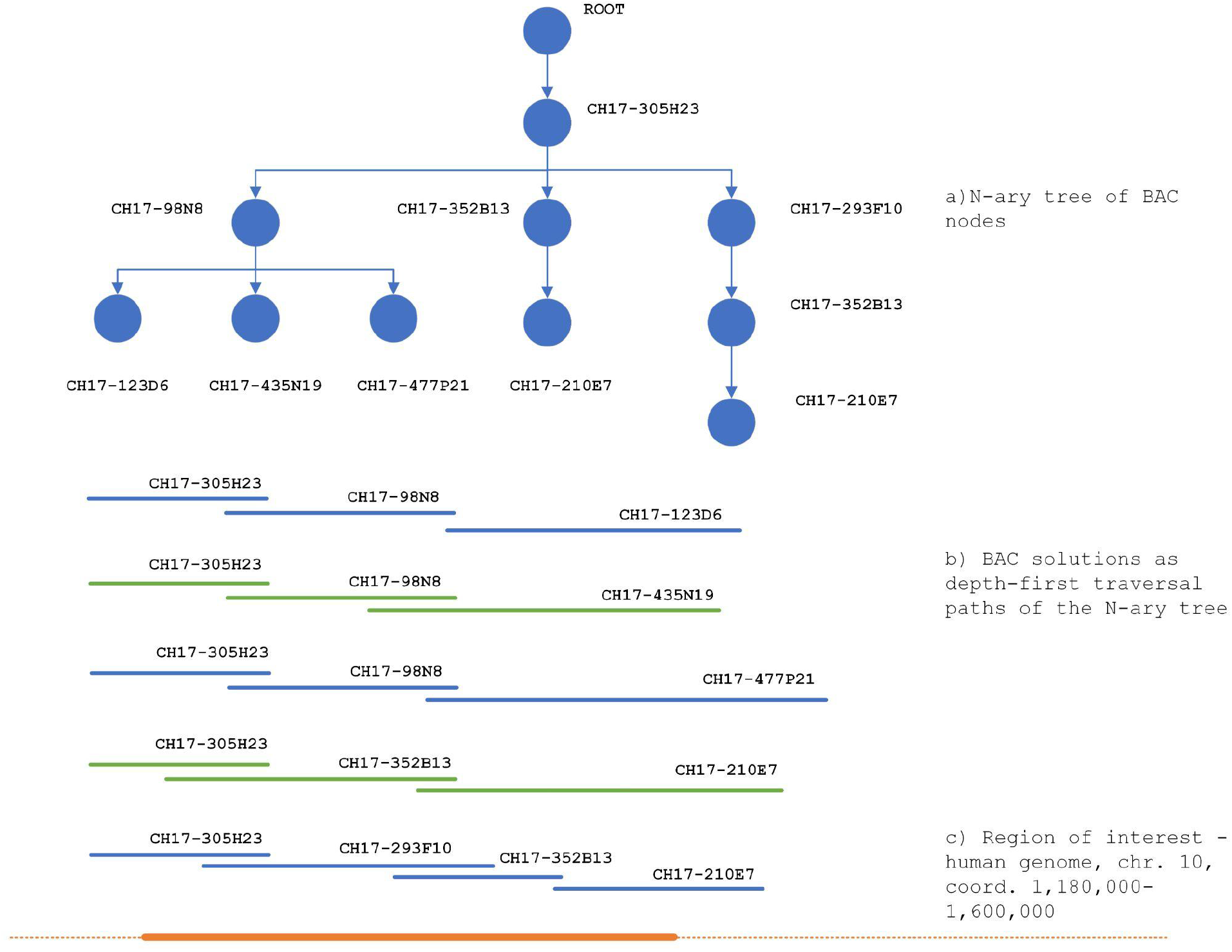
A portion of a BAC N-ary tree for a region in the human genome hg38, chromosome 10, coord 1,180,000 - 1,600,000. Each depth-first traversal of the tree constitutes a BAC solution for this region.

We have designed a recursive algorithm for parsing BAC libraries in search of the BAC solutions for a given region of interest. By applying this algorithm, we build our N-ary tree of such BAC solutions. Here, the main property of this tree is the fact that each depth first traversal of a tree represents a unique BAC solution for the region of interest. A complete list of these solutions could be obtained as a collection of all depth first traversal paths for the tree. Java implementation of the algorithm is available on the MendelProject gitHub site.

To ensure efficient parsing of BAC libraries we have ported libraries for supported genomes to the internal database tables. Currently, BAC libraries from human, mouse and other species are available/supported. To further speed up the search process, We have indexed these tables on key columns - library name, chromosome, start location.

A class diagram for the concrete implementation of MenDEL’s BAC finder is presented in Fig. 3. Here, BACNodeTree class extends abstract NaryTree and NaryNode classes while implementing axillary interfaces such as Rangeable, Jsonable, Markable. The NaryPath template class represents a depth-first traversal of NaryTree. When parametrized by the BACNode class, an object of the NaryPath represents a BAC solution.

**Figure 3.**
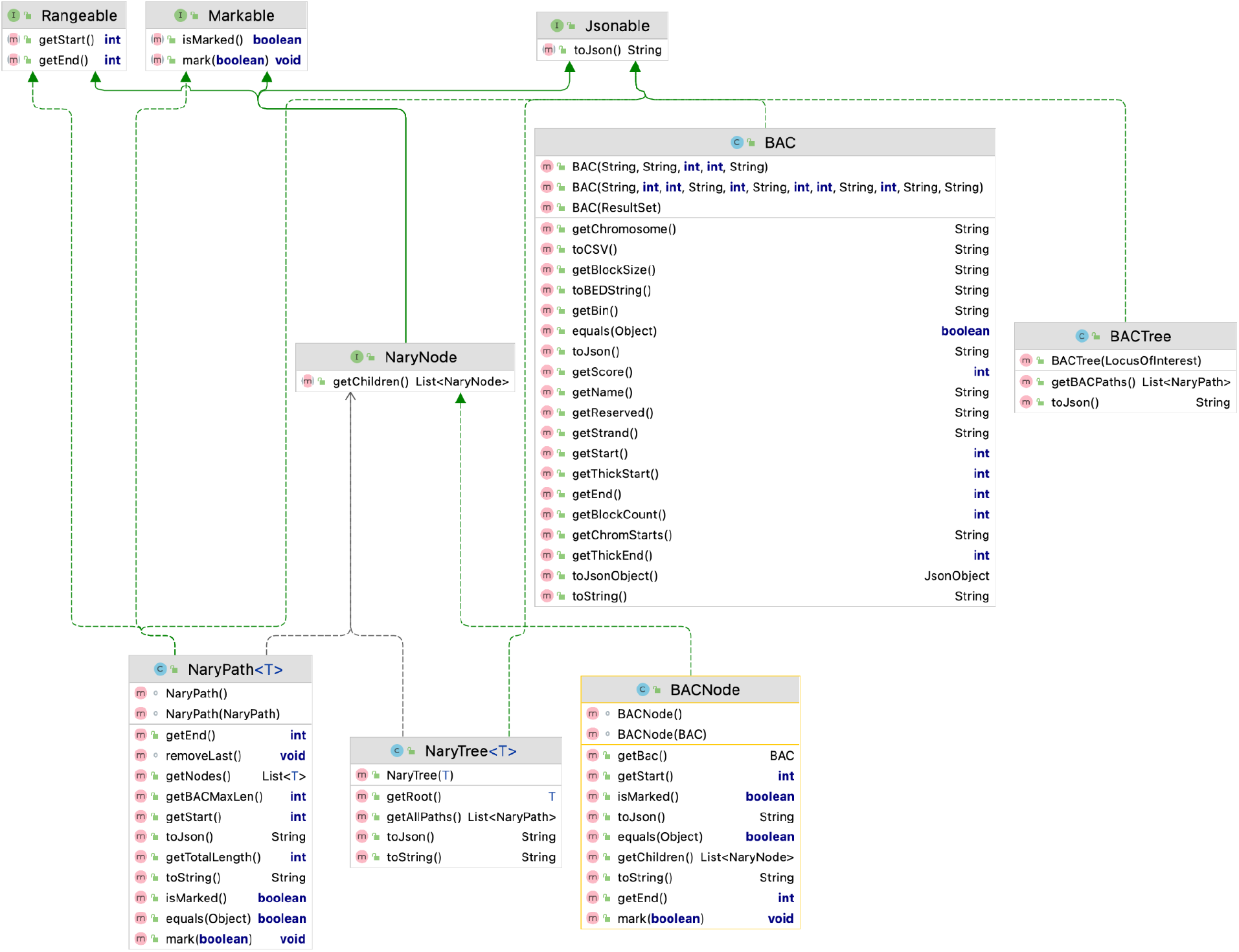
Class diagram for MenDEL’s BAC finder implementation

An immutable BACTree object is constructed by calling a constructor for the specified region of interest. Then, the BAC solutions are returned as a collection to a user or a client application by calling the getBACPaths() method of the BACTree object. This collection of BAC solutions could be sorted based on various parameters - length, number of BAC nodes, etc.

### 2.2 MenDEL - BAC Finder module

The BAC Finder classes are used as components in a number of MenDEL modules. They also constitute the core of the BAC Finder module. The user interface for the BAC Finder module is shown in Fig.4.

**Figure 4.**
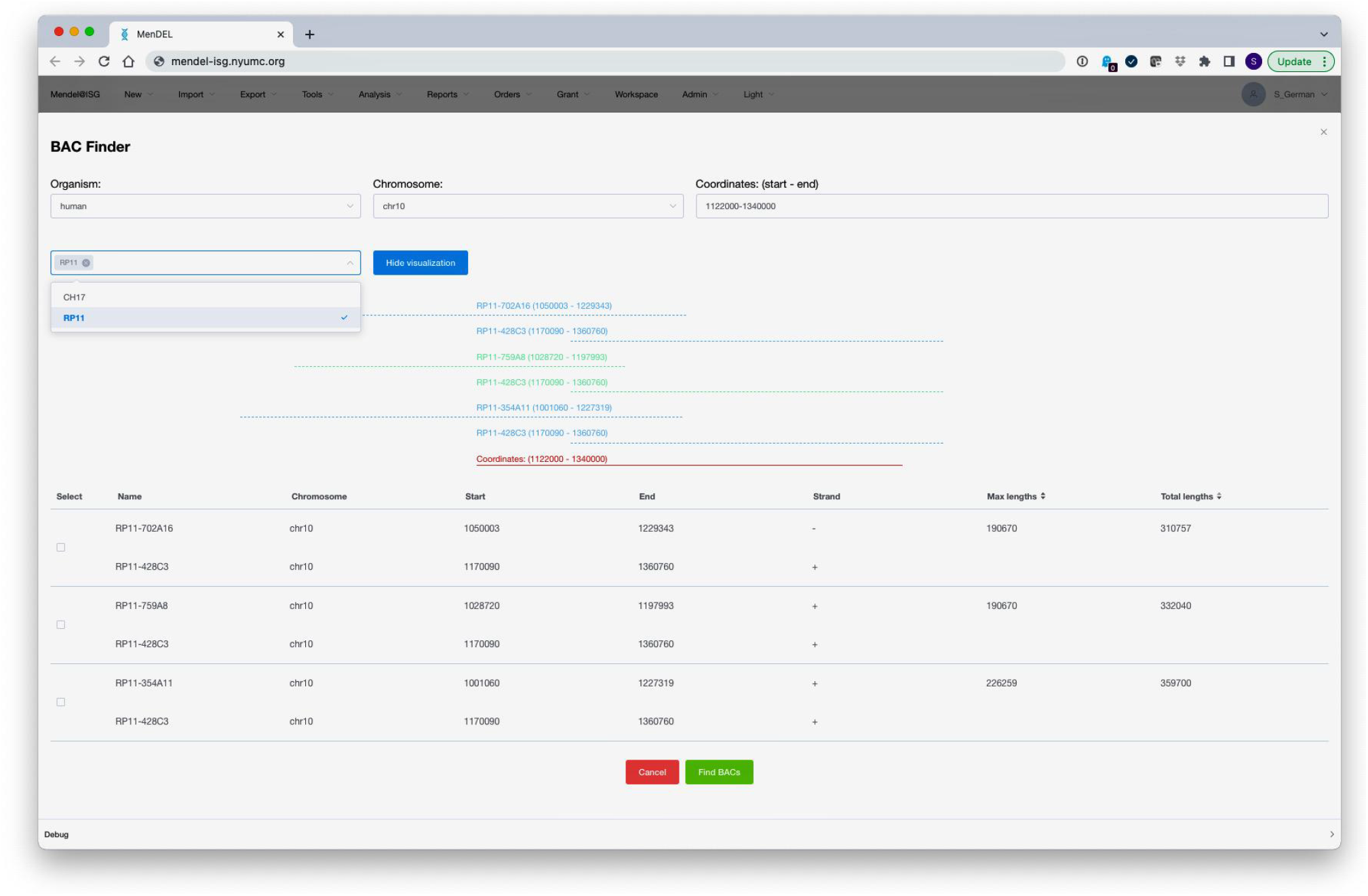
The user interface for the BAC Finder Module.

In this module users can specify their region of interest by choosing an organism, a chromosome, and start and end coordinates for the region. Users can further filter results by a library (CH17 or RP11 in Fig. 2) Results can be sorted based on the total solution length, the length of the longest BAC, number of BACs. The results are visualized for user convenience.

## 3. Conclusion

The interpretation of the BAC solutions as a collection of depth-first traversal of N-ary tree, where nodes are associated with BAC objects provides compact and efficient data structures and algorithms for automated search for such solutions for a given region of interest. We build these N-ary trees by utilizing recursive algorithms for parsing BAC libraries. Deploying BAC libraries as indexed database tables allows further speed up and automate parsing of these libraries. An important property of those N-ary trees is that their depth-first traversals provide a complete list of unique BAC solutions. Each of these BAC solutions is an object, and collection of these objects could be sorted and filtered by a number of parameters - source library, total length, longest BAC, etc. Finding, sorting, and filtering BAC solutions for reasonably long regions of interest without automation would be a time consuming and error-prone process.

Efficiency of parsing and search algorithms, flexibility of filtering and sorting of results, ease of use, and possibility of customization makes interpretation of BAC solutions as N-ary trees a useful approach in designing non-trivial DNA assemblies.

## Acknowledgments

This work was supported in part by NIH/NHGRI grant 1RM1HG009491.

